# Phylogenetic diversity and functional potential of large and cell-associated viruses in the Bay of Bengal

**DOI:** 10.1101/2023.07.17.548743

**Authors:** Benjamin Minch, Salma Akter, Alaina Weinheimer, M Shaminur Rahman, Md Anowar Khasru Parvez, Sabita Rezwana Rahman, Md Firoz Ahmed, Mohammad Moniruzzaman

**Affiliations:** Department of Marine Biology and Ecology, Rosenstiel School of Marine, Atmospheric, and Earth Sciences, University of Miami, Miami, FL 33149; Department of Microbiology, Jahangirnagar University, Savar, Dhaka-1342, Bangladesh; Bigelow Laboratory for Ocean Science, 60 Bigelow Drive, East Boothbay, ME 04544; Department of Microbiology, Jashore University of Science and Technology, Jashore, Bangladesh; Department of Microbiology, University of Dhaka, Dhaka-1000, Bangladesh

## Abstract

The Bay of Bengal (BoB), the largest bay in the world, provides valuable ecosystem services such as fishing and recreation to millions of people living along its coast and has a significant economic value. The BoB is impacted by various environmental factors such as seasonal monsoons and multiple freshwater inputs, and this region is particularly vulnerable to sea-level rise and increased frequency of devastating cyclones that are predicted to be exacerbated due to global climate change. These factors are also compounded by anthropogenic influences from tourism and development, making it an important ecosystem to understand and study from a global change perspective. Despite its importance, microbial diversity and ecology have remained largely understudied in this region. In this study, we describe the diversity and putative functional importance of large and cell-associated (that is, originating from the cellular size fraction) viruses from two coastal sites in the BoB, with an emphasis on giant viruses and large phages. Sites chosen for this study include Cox’s Bazar, a populated beach with multiple freshwater inputs, and Saint Martin Island, a resort island that has considerably less human influence compared to Cox’s Bazar. Through metagenomic sequencing, we were able to identify a more abundant and more diverse viral community at Cox’s Bazar consisting of many viruses that are indicators of freshwater intrusion and runoff. Overall, 1962 putative phage genome bins were obtained ranging from 10 - 655 kilobase pairs (kbp) in sizes. Of these genomes, 16 from Saint Martin were found to be larger than 100kbp which we deemed “large” phages, and we were able to reconstruct a phylogeny of these large phages using the TerL gene as a marker. This phylogeny revealed clades enriched in large phages and a high diversity of large phage candidates in the Bay of Bengal coast. Protein annotation analysis showed a wide variety of functionality from both sites with more auxiliary metabolic genes (AMGs) found in the Cox’s Bazar viral community. Five giant virus (Phylum Nucleocytoviricota) genomes were also reconstructed from Cox’s Bazar and identified as belonging to the orders Imitervirales and Pandoravirales. These genomes ranged from 83 - 876 kbp in size and contained a wide range of encoded functionalities. To the best of our knowledge, our study represents the first insights on the phylogenetic and functional diversity of viruses in the Bay of Bengal. These results thus provide an important foundation for further studies on the impact of host-virus interactions on biogeochemical cycles and microbial food web in this understudied marine environment.

## Introduction

The Bay of Bengal (BoB) is the largest bay in the world and provides ecological and economic services to all surrounding nations. Home to many unique habitats such as seagrass beds, coral reefs, and mangrove forests, the BoB is an important biodiversity hotspot. The diversity of habitats and species in BoB has made it an important ecotourism destination in surrounding nations, with many countries developing their tourism economies around activities in the bay. In addition to ecotourism, the BoB is also a source of many natural resources. One such resource is the many fisheries along the coast of the BoB. These fisheries have been described as some of the most productive in the world, given the wide range of species and the vast area of water in the bay (Islam, 2003). In Bangladesh, these fisheries are the second largest source of employment, making up 8% of the workforce and employing around 13 million people (DOF, 2010; Hossain & Islam., 2006). These fisheries are supported by a large population of coastal mangrove habitats as well as a diverse assemblage of microbes making up the bottom of the food web.

Despite its importance, the Bay of Bengal is one of the most understudied bodies of water in the world (Masud-Ul-Alam, 2020). Consequently, the microbial communities at the foundation of the BoB ecosystem remain poorly understood. A few recent studies have looked at bacterial diversity through 16S amplicon sequencing, but very few have looked at the functional potential and interactions of the microbial communities in this region (Rajpathak et al., 2018, Wu et al., 2022). Among the studies that have attempted to look at microbial diversity through metagenomics, all have been focused on sediment bacteria in the deep sea (Marimuthu et al., 2022), with little or no efforts targeted towards understanding the microbiota of surface water communities, the communities closely associated with corals, seagrass, and mangroves using similar approach. Unlike taxonomic-focused studies that exclusively enumerate which species are present, metagenomic studies provide functional information about these microbial communities related exclusively to their role in the broader biogeochemical cycling of the ecosystems.

Another substantial gap in our understanding of the microbial dynamics of BoB is that all previous studies on the microbial community have excluded viruses. The one exception to this is the study of the white spot syndrome virus, a virus infecting black tiger shrimp, important aquaculture species in the BoB (Debnath et al., 2014). Viruses are the most abundant biological entities in the ocean and influence nutrient cycling, biogeochemical cycles, population dynamics of prokaryotes and microeukaryotes, algal bloom dynamics, and numerous other ecological processes (Fuhrman, 1999). It is estimated that there are about 10^9^ viruses per liter of water and that they are responsible for the death of roughly 10 to 50% of the bacteria in the surface ocean every day (Fuhrman, 1999, Suttle, 2007). In addition to simply killing their host, viruses have been shown to modulate host metabolism through the use of auxiliary metabolic genes (AMGs) (Luo et al., 2022). These AMGs have large implications for molecular biogeochemical cycling as they have the potential to alter host nutrient uptake, photosynthesis, and carbon metabolism (Heyerhoff et al., 2022).

While most viruses likely contain AMGs, certain viruses have much larger genomes and potentially harbor more metabolic capacities to modulate host metabolism. Jumbo phages, bacteriophages with large genomes and large capsids, are prime examples of this as their large genomes can potentially contain more AMGs than an average smaller bacteriophage that has fewer genes (Hendrix, 2009). These jumbo phages have until recently been underrepresented in virome data due to their large size and filtering strategies commonly adopted in environmental virome studies (Al-Shayeb et al., 2020; Weinheimer et al., 2020). Another group of large viruses is the Nucleocytoplasmic Large DNA Viruses (NCLDV). These DNA viruses mainly infect microbial eukaryotes (also known as protists), typically have large genomes (upwards of 2.5Mb) and are widespread in the Earth’s oceans (Schulz et al., 2022; Moniruzzaman et al., 2020). NCLDVs have received recent attention due to their wide genomic repertoire and potential for metabolic reconstruction in their host (Moniruzzaman et al., 2020; Ha et al, 2021). Both jumbo phages and NCLDVs present a huge potential for housing a large number of AMGs and thus have a substantial effect on host metabolism and biogeochemical cycles.

Viruses, along with other members of the microbial community have been shown to shift their dynamics and diversity in response to various environmental and anthropogenic stressors (Bisset et al., 2013; Danovaro et al., 2003). These stressors are abundant in the Bay of Bengal as many of the bordering countries are developing at fast rates and still lack basic environmental regulations present in more developed nations. For example, the regulation of wastewater remains a large problem for coastal communities along the bay. With large populations and underdeveloped wastewater treatment plants, much of the household and industrial wastewater makes its way into rivers and the bay (Asaduzzaman, 2020). Additionally, countries surrounding the BoB are hit by frequent monsoons which lead to agricultural runoff of fertilizers and pesticides (Sinha et al., 2022). This runoff, along with other environmental stressors, has the potential to shift the microbial community as nutrients and waste products are added to the waterways.

Furthermore, understanding the effects of environmental stressors on microbial communities in the Bay of Bengal requires an examination of the viral diversity and its association with host organisms. Despite the potential impact of runoff water on microbial population dynamics, the viral community in this region and its interactions with hosts have yet to be characterized. Addressing this gap, our study reports the composition and dynamics of the viral community at two distinct sites on the eastern side of the Bay of Bengal, Bangladesh. Specifically, using metagenomic data, we sought to identify the large and cell-associated viruses (viruses present in the cellular size fraction) present at these sites to characterize the diversity and abundance of viral populations in this under-studied ecosystem. Functional annotations and host predictions were also performed to examine the roles these viruses could be playing in the metabolism of the respective microbial communities. To our knowledge, this is the first characterization of the cell-associated viral populations in this region. This foundational work is crucial to inform future investigation and ecological characterization of this unique ecosystem and its interplay with the developing human civilizations around it.

## Methods

### Sample collection, processing and sequencing

Samples were collected from 4 different sites at both Saint Martin and Cox’s Bazar, Bangladesh. These four sites were aggregated into a single sample for each location which from this point forward will be referred to as S1 for Saint Martin and S2 for Cox’s Bazar. These were collected from seawater between March 2 and 3, 2022 during low tide. Samples were collected in 1 L sterile sampling bottles at 1.5m depth from the surface. After collection, samples were processed at the Microbiology laboratory at Jahangirnagar University, Savar, Dhaka, Bangladesh. The geographic location of sampling, as well as the measured environmental parameters, are shown in Figure 1.

**Figure 1.**
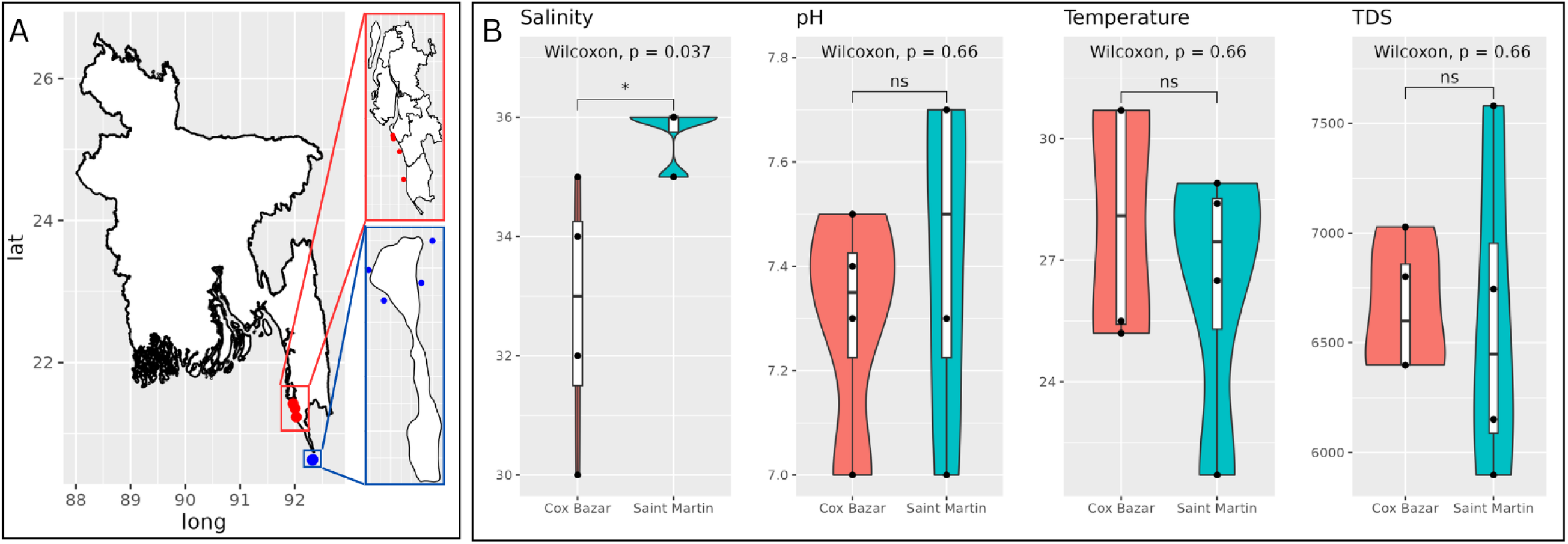
Sampling Location and physicochemical parameters. (**A**) A map of Bangladesh and study locations. Cox’s Bazar is highlighted in red, and Saint Martin is highlighted in blue. (B) Physicochemical measurements of the two sites. Violin plots represent the measurements from 4 locations at each site with a boxplot in the center showing the mean and range of the data. Sites were compared using a Wilcoxon test.

Water samples were first passed through Whatman filter paper no. 1 (pore size 11um) and then subsequently passed through a 0.45um membrane and 0.2um membrane. From each of these two filters, DNA was extracted using the DNeasy PowerWater Kit (QIAGEN) according to the manufacturer’s protocol. Purified DNA samples from each membrane were mixed together and sent to EzBiome Inc., USA for metagenomic sequencing. An equal quantity of DNA from both membranes from the four representative sites were pooled together as a single sample from each site.

Paired-end (2x 150bp) sequencing was performed using a Novaseq 6000 sequencer (Illumina Inc., USA). The FASTQ files were evaluated for quality using FASTQC (v0.11) (Andrews, 2010) and adapter sequences and low-quality ends were trimmed using Trimmomatic (v0.39) (Bolger et al., 2014). After trimming, the read counts for S1 and S2 were 33.94 and 31.8 million respectively, corresponding to 92.2 and 92.37% of total reads.

### Identification and characterization of the phage community

Metagenomic reads from both sites were trimmed using Cutadapt (v4.4) (Martin,2011) and then assembled using MetaSpades (v3.15.5) (Prjibelski et al., 2020). These assemblies from S1 and S2 were then analyzed using the ViWrap pipeline (v1.2.1) with default settings to bin viral contigs as well as classify them and predict their functional capacities (Zhou et al.,2023). ViWrap uses a combination of Virsorter2 (v2.4.5), VIBRANT (v1.2.1), and DeepVirFinder (v2020.11.21) to identify viral contigs. Contigs are then binned using vRhyme (v1.1.0). Phages are then classified using the VOG HMM database (VOG 97) and NCBI RefSeq (release 218) viral protein database. Host prediction was also performed in this pipeline using the iPHOP module (v1.2.0).

Viral contigs obtained from ViWrap were de-replicated using dREP (v3.4.0) (Olm et al., 2017) with a 95% ANI cutoff and quality assessed with CheckV, resulting in a total of 1962 viral sequences (a mix of binned and unbinned contigs) that had a CheckV quality assignment of medium or higher. This set of de-replicated sequences was used for read mapping from both sites. Read mapping was done using minimap2 (v2.24) (Li, 2018) and then coverage was determined using CoverM (in ‘genome’ mode) with a minimum identity of 95% (v0.6.1) (Robbins et al., 2017). Graphs of viral abundance using RPKM (reads per kilobase per million) were generated using ggplot2 in R.

### Phylogenetic analysis

For MCP (major capsid protein) phylogenetic analysis of NCLDV, MCP proteins belonging to NCLDV members were identified using NCLDV markersearch script (Moniruzzaman et al., 2020). This set of MCP proteins was then de-replicated at 95% using cd-hit (v4.8.1) (Fu et al., 2012). Sequences from MCP were aligned to a reference set of MCP proteins from known NCLDV derived from the Giant Virus Database (GVDB) (Aylward et al., 2021). MAFFT was used for alignment (v7.511) (Katoh et al., 2013) using default parameters. Aligned sequences were then made into a maximum likelihood tree using fasttree with default settings (v2.1) (Price et al., 2010). The tree was visualized using iTol (Letunic & Bork,2007).

Phylogeny of the TerL (terminase large subunit) gene found in bacteriophages was reconstructed using sequences derived from our data as well as multiple different databases. Reference sequences were obtained from the VOG database (http://vogdb.org/), NCBI Refseq database (O’Leary et al., 2016), the infrared database (Cook et al, 2021), and data from Al-Shayeb et al (2020), and Weinheimer et al (2022). Metadata on genome size, location of isolation, and identification can be found in the supplemental information (see Data Availability statement). TerL sequences from our dataset were identified using HMMER3 (v3.3.2) (Eddy, 2011) with an E-value cutoff of 1e-5 against all the TerL HMM profiles from the VOG database. All of these sequences were clustered at 95% ANI using cd-hit and aligned with MAFFT using the auto setting. The alignment was then trimmed using TrimAL (v1.3) (Capella-Gutiérrez et al., 2009) with parameter ‘-gt 0.1’ and a phylogenetic tree was reconstructed using IQ-Tree (v2.2) (Nguyen et al., 2015) with a LG+F+R10 model. The tree was visualized using iTol (Letunic & Bork,2007). A tree with the bootstrap values is available in the supplemental data provided.

### NCLDV Genome Binning and Contig Prediction

A standardized pipeline was used to identify and bin NCLDV genomes in our dataset. For NCLDV functional analysis, assembled contigs were binned using metabat2 (v2.12.1) (Kang et al., 2019). Proteins were predicted for the bins using prodigal-gv (v2.11) (Hyatt et al., 2010), and then bins were screened for the presence of NCLDV marker genes using the NCLDV markersearch script. Bins with at least one hit to an NCLDV marker gene (MCP, SFII, RNAPS, RNAPL, PolB, TFIIB, TopoII, A32, VLTF3) were kept and screened for signatures of NCLDV contigs using ViralRecall (Aylward & Moniruzzaman, 2021). Bins with a positive ViralRecall score were kept for further screening which involved the removal of bins not fitting the criteria of having 3 of the 4 key marker genes described in Aylward et al (2019). After screening, contigs with negative ViralRecall scores were removed from the bins as they most likely represent bacterial or eukaryote contamination. Following this stringent protocol resulted in 5 NCLDV genomes. tRNAs for these genomes were predicted with tRNAscan-SE (Lowe & Chan, 2016).

### Functional Analysis

Functional annotations of the genes in the NCLDV genomes were assigned using HMMER3 against the GVOG (Aylward et al., 2021), Pfam (Finn et al., 2014), and EGGNOG (Cantalapiedra et al., 2021) databases using an E-value cutoff of 1e-5. Viral contigs that were not binned into a genome were also identified with ViralRecall and annotated in a similar fashion with HMMER3. In total, there were 60 contigs found in S2 and 39 found in S1 that were not present within the NCLDV MAGs. Functional categories were then graphed as a proportion of total functionality using ggplot2. Maps of giant virus genomes were also made with the circlizewith circlize (v0.4.15) (Gu et al., 2014) package in R.

For prediction of gene functionalities of phages, the ViWrap pipeline was used, which employs VIBRANT to categorize AMGs (auxiliary metabolic genes) of viral contigs. After running ViWrap for each site, AMGs coming from large phages (those with genomes or unbinned contig sizes over 100kbp) were separated out for comparison. Resulting functional plots were made using the ggplot2 package in R.

## Results

### Diversity, Abundance, and Functional Potential of Prokaryotic Viruses

We obtained a total of 1962 phage viral genomes from both Saint Martin’s Island and Cox’s Bazar that passed the screening criteria of ViWrap (see methods). Of these, 1058 (53.92%) were unique to Cox’s Bazar and 843 (42.97%) were unique to Saint Martin at a 95% Average Nucleotide Identity (ANI) cutoff. 61 genomes were shared between sites (3.01%).

A comparison of alpha diversity between Saint. Martin and Cox’s Bazar revealed Cox’s Bazar to have both a higher Shannon diversity and relative abundance of viruses with mean log RPKM of 3 compared to 2.6 for Saint Martin (Figure 2A-B). Many viruses exhibited site-specific abundance patterns and were categorized into three groups: those exclusive to Saint Martin, those exclusive to Cox’s Bazar, and those found at both sites. The dominant viruses at Saint Martin showed homology to phages that belong to species that infect *Puniceispirillum*, *Synechococcus*, *Vibrio*, *Pelagibacter*, and *Lentibacter*. Meanwhile, the phages unique to Cox’s Bazar were mainly composed of phages with homology to species that infect *Synechococcus*, *Flavobacterium*, *Cyanobacteria*, *Puniceispillium*, and *Pelagibacter* (Figure 2C).

**Figure 2.**
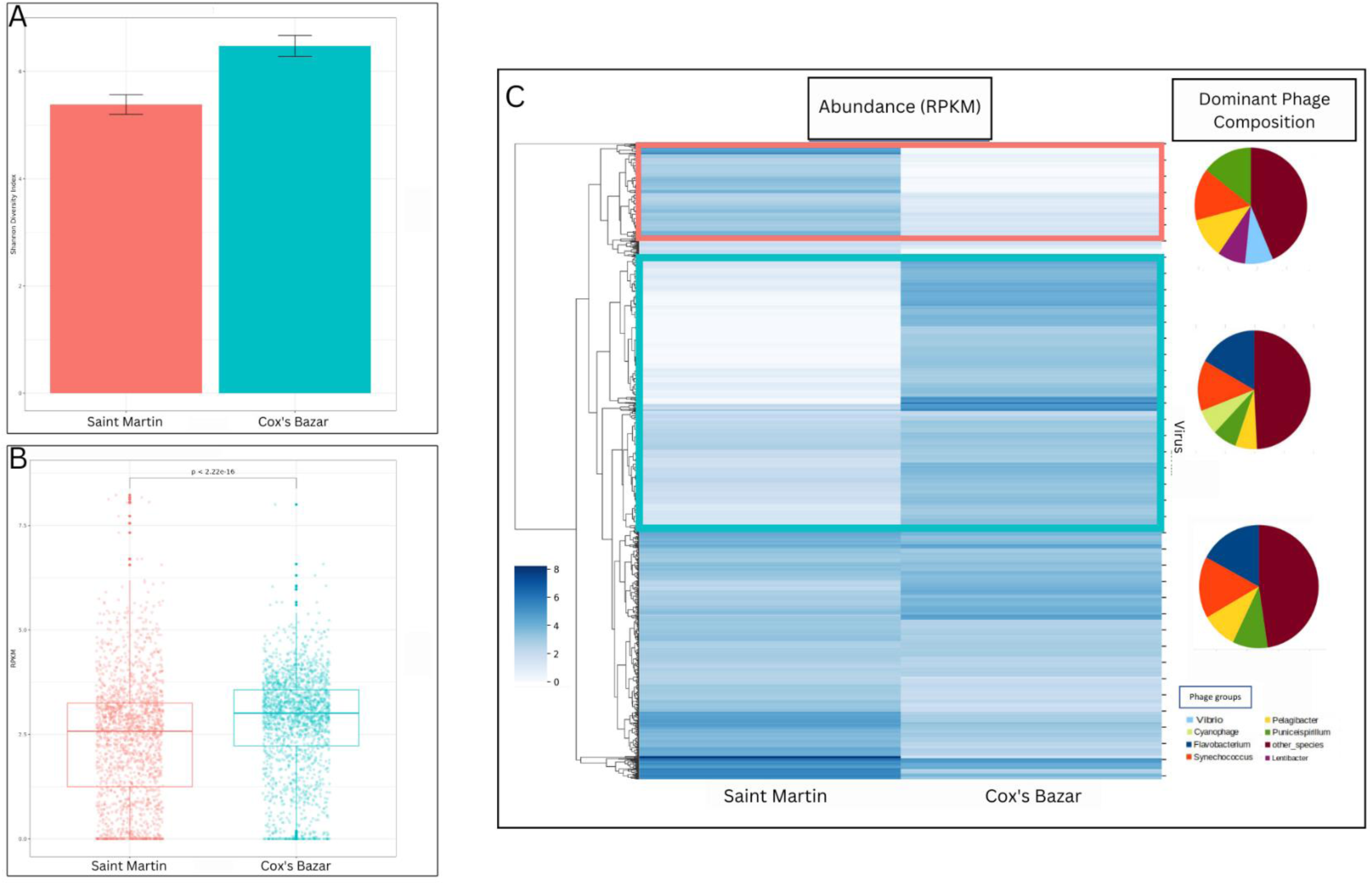
Prokaryotic Viral Abundance and Diversity. **(A)** Shannon diversity indices were calculated from viral populations at both sites. Error bars represent +/- 2 SEM. **(B)** Comparison of the overall abundance of prokaryotic viruses at each site. Each dot represents a virus and its measured abundance (Reads per kilobase per million [RPKM]). A t-test was performed on the mean RPKM between sites (p <2.22e-16). **(C)** Heatmap showing viral abundance and dominant phage composition of each cluster. Each line in the heatmap represents one virus and its abundance between the two sites. Three clusters were formed from this heatmap: viruses present almost exclusively in Saint Martin (red box), viruses present almost exclusively in Cox’s Bazar (blue box), and viruses present in both sites (no box). The dominant phage groups are shown for each of these three clusters, represented as a proportion of the total cluster population.

In both sites, most of the DNA viruses were prokaryotic, belonging to the prokaryotic virus class of Caudoviricetes, followed by the giant virus class Megaviricetes. In general, the abundance of different viral classes was similar between sites with the exception of the virophage class Maveriviricetes as more prevalent in Cox’s Bazar and the Tectiliviricetes class, which includes prokaryotic viruses and adenoviruses of vertebrates, as more prevalent in Saint Martin (Figure 3A).

**Figure 3.**
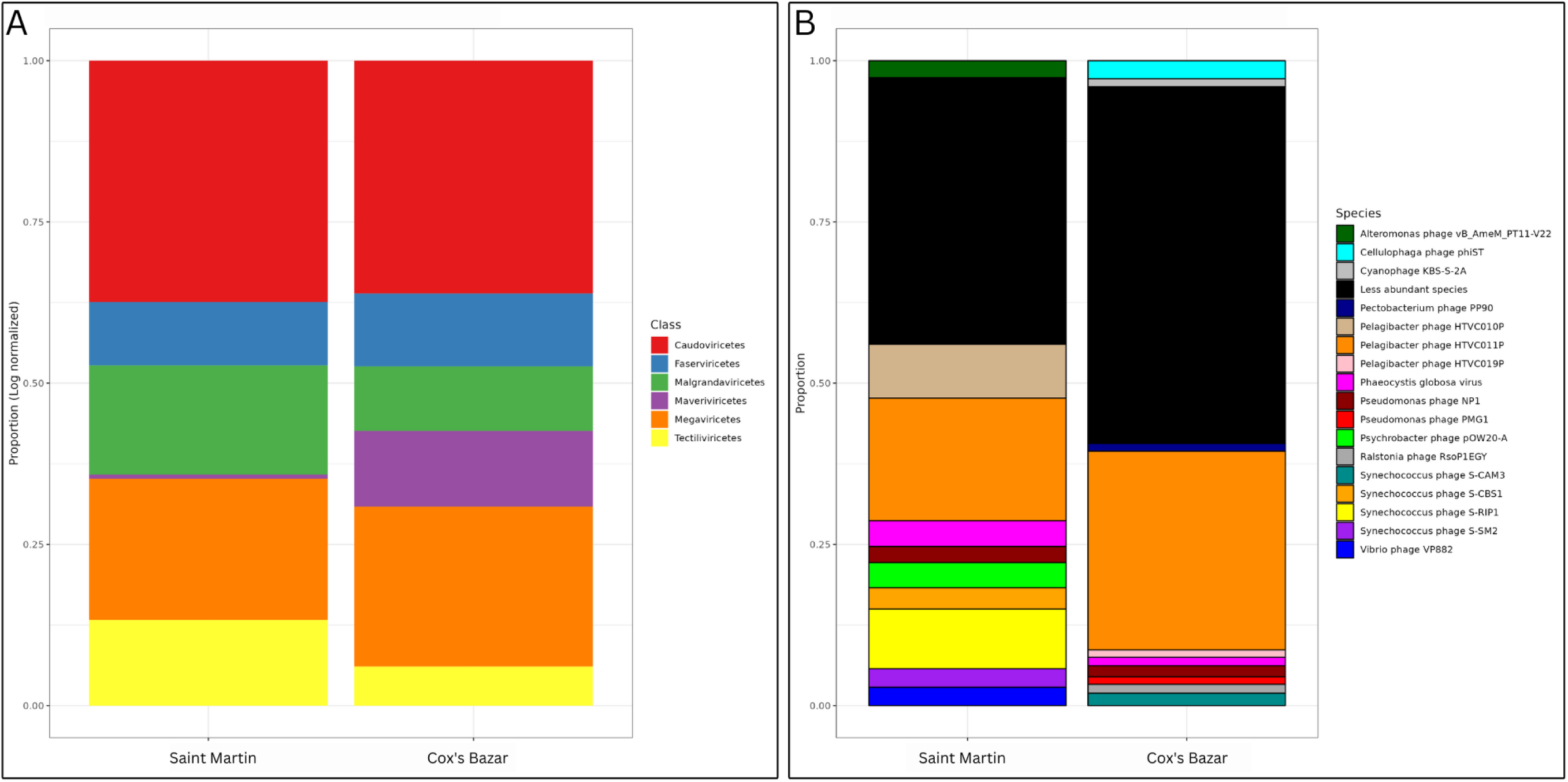
Viral Community Composition. **(A)** Class-level and **(B)** Species-level taxonomic distribution of viruses between sites. Proportions are log normalized for class-level taxonomy to see better resolution. For the species-level taxonomy, only the top ten most abundant species from each site are shown as a unique bar on the graph. All other species are grouped into the black bar titled ‘less abundant species. Taxonomic assignments were obtained through the ViWrap pipeline.

When looking at the top 10 most abundant species (that were taxonomically identifiable) of phages in each site, there are some clear differences. Most of the phages in Saint Martin belonged to these top species, while the community of Cox’s Bazar was dominated by species that were rarer. The dominant viral species in both communities, however, is the *Pelagibacter* phage HTVCO11P. This is consistent with the fact that this species is known to infect members of the *Pelagibacter* genus, one of the most abundant bacterial genera in the ocean (Giovannoni, 2017). Interestingly, Saint Martin hosts a larger proportion of phages belonging to the species *Psychrobacter* phage pOW20-A, than Cox’s Bazar. Although this phage species is normally found in cold Arctic or Antarctic waters, a previous study detected the bacterial genera Psychrobacter as dominating the coastal regions of the BoB (Wu et al., 2022), serving as a potential host for members of this Psychrobacter phage pOW20-A species. Saint Martin also has a larger proportion of phages with homology to species of *Vibrio* and *Alteromonas* phages than Cox’s Bazar (Figure 3B). The study by Wu et al (2022) found bacteria of these phage’s putative host genera to be more prevalent in offshore waters.

The species composition of the phages indicates that Cox’s Bazar has influences from its nearness to freshwater and potential sewage inputs. The *Cellulophaga* phage phiST species was one of the top phage species in Cox’s Bazar, but not dominant in Saint Martin. The marine bacteria *Cellulophaga* typically increases in abundance during high nutrient and bloom-forming conditions, as they are responsible for the degradation of biopolymers like polysaccharides, and thus are involved in recycling the organic matter derived from blooms (Bischof et al., 2021), making it unsurprising to find a prevalence of Cellulophaga phage species in Cox’s Bazar. *Ralstonia* phage was also uniquely dominant in Cox’s Bazar and suggest runoff input into this site since the bacteria of the genera *Ralstonia* are usually found in freshwater and soil environments often as pathogens for crops. The presence of this phage species could be indicative of runoff and freshwater inputs near the BoB (Vogelaar et al., 2023). Cyanophages were also found in much higher abundance at Cox’s Bazar, but it is unclear how this enrichment relates to terrestrial proximity compared to Saint Martin since cyanophages are present in both freshwater and marine habitats (Xia et al, 2013).

Overall, 250 phage-encoded auxiliary metabolic genes (AMGs) were identified in phages from Saint Martin and 524 AMGs were identified from Cox’s Bazar (Figure 4). Of these, 100 belonged to the genomes of large phages (> 100kbp). Most of these AMGs were categorized as being involved in the metabolism of cofactors and vitamins - with the most abundant pathways being the biosynthesis of folate and the metabolism of porphyrin and chlorophyll. A large number of AMGs putatively involved in cysteine and methionine metabolism, lipopolysaccharide biosynthesis, and amino sugar and nucleotide sugar metabolism were also found in these genomes. Cox’s Bazar had 15 more pathways that weren’t detected in Saint Martin’s Island - which included pathways involved in methane metabolism, glycolysis, the TCA cycle, carbon fixation in photosynthetic organisms, pyruvate metabolism, glycine, serine, and threonine metabolism. Phosphonate metabolism and glycosphingolipid biosynthesis were the only pathways found to be unique to samples from Saint Martin.

**Figure 4.**
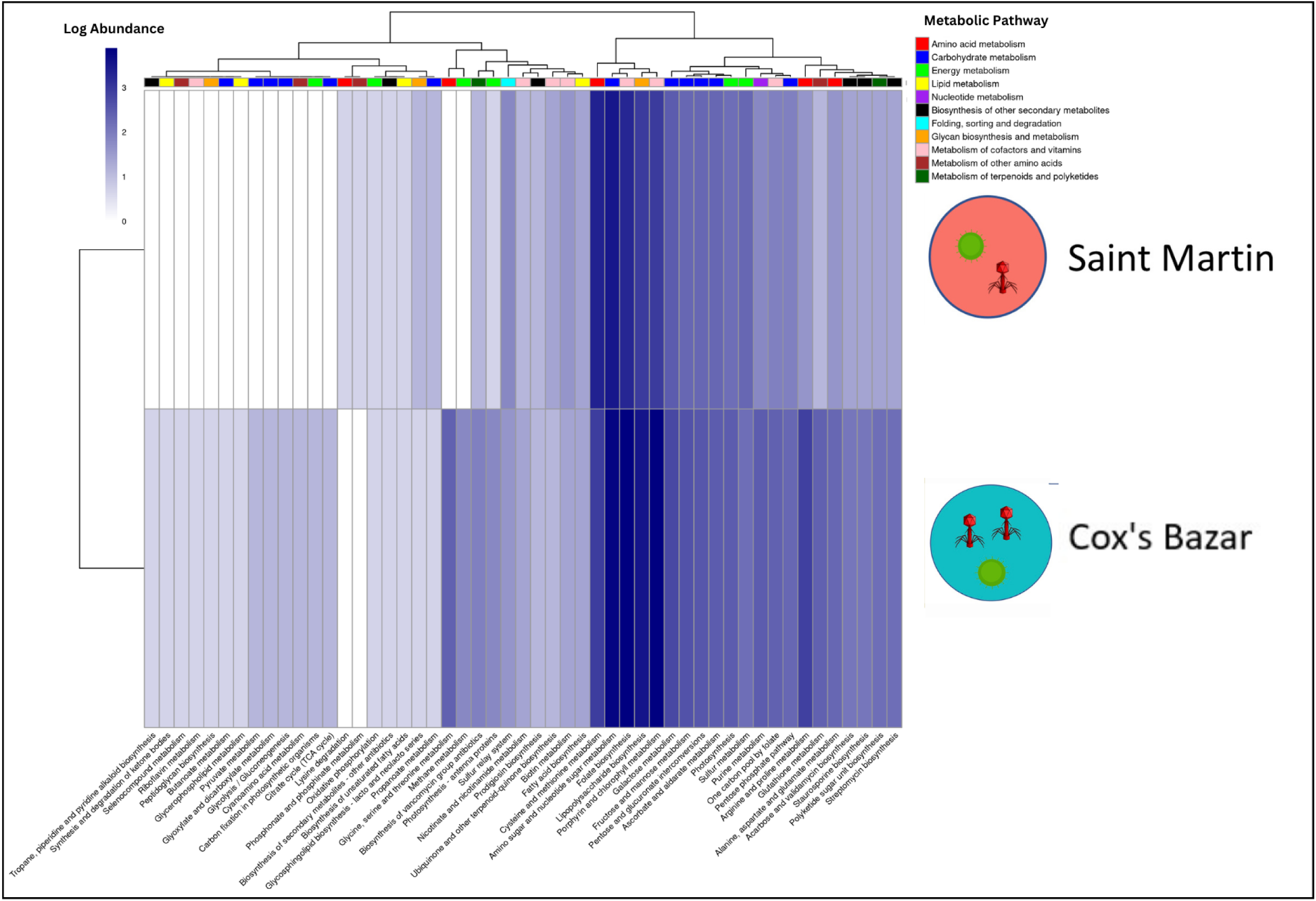
Functional Profiling of Prokaryotic Viruses. A heatmap showing VIBRANT-predicted auxiliary metabolic genes (AMGs) of prokaryotic viruses from both locations. Values shown are log abundance of AMGs in each metabolic pathway, with larger metabolic categories being shown in the color strip at the top of the graph.

### Large Phage Diversity and Functional Potential

The nomenclature for large phages is not yet definitive, likely because the size cutoffs used to define these groups have no evolutionary basis and these phages are likely paraphyletic, having evolved from smaller phages in multiple, independent occasions. For instance, phages with genomes over 200 kilobases have traditionally been called “jumbo phages”, but a recent study enumerating jumbo phages in diverse metagenomes referred to them as “huge phages” and created the term “megaphage” to refer to those with genomes over 500 kilobases. Although we recognize that most papers focused on jumbo phages refer to the 200kbp cutoff (Hendrix, 2009; Weinheimer et al., 2022), for this study, we have opted to include all phages over 100kbp in size in order to capture a broader range of these phages. For this work, we were primarily interested in identifying jumbo phages based on their phylogenies, and this cutoff was chosen to evaluate functional differences between large and smaller phages. Using this criterion, we identified 16 phages with genome sizes larger than 100kbp from Cox’s Bazar, ranging from (102-655kbp).

From our phylogenetic analysis of the TerL (terminase large subunit) gene, there emerged clades enriched with known large phages that cluster closely on the tree (Figure 5A). Many of our own phage TerL genes from Cox’s Bazar and Saint Martin also fell inside these clades, confirming the presence of jumbo phages in our sites and adding some support to the chosen 100kbp cutoff. The remaining jumbo phages in our dataset cluster close to other known jumbo phages on the tree, although not clustering inside of the enriched clades. There was generally no trend when it comes to site-specific clustering or isolation origin although the jumbo clade is enriched in marine viruses with many non-marine jumbos clustering elsewhere.

**Figure 5.**
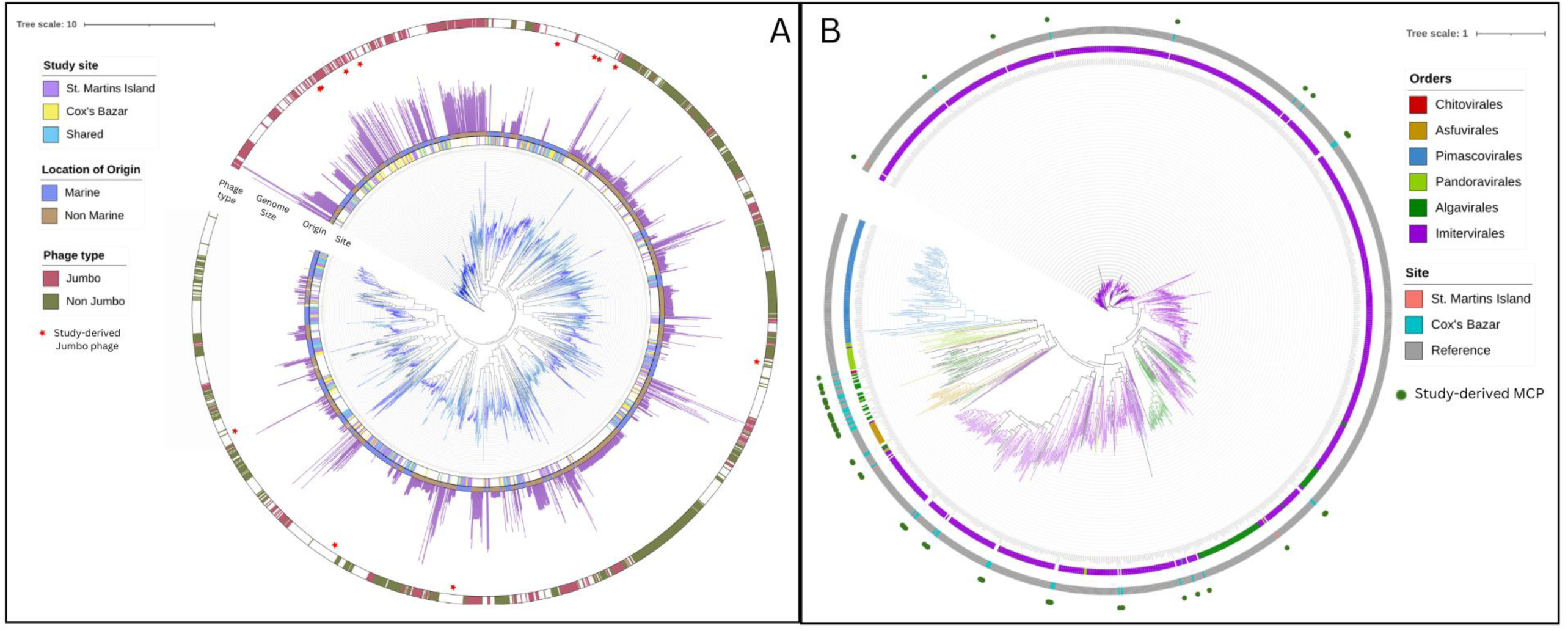
Phylogenetic analysis of large viruses. **(A)** Large phage phylogeny was constructed using TerL (terminase large subunit) as a marker gene with the LG+F+R10 model in IQ-TREE. The tree includes all phages with a TerL gene from our study as well as reference sequences from the VOG, NCBI Refseq, and the infrared database, as well as known jumbo phages from Al-shayeb et al (2020), and Weinheimer et al (2022). **(B)** Phylogeny of the NCLDVs found in this study using the major capsid protein (MCP) as a phylogenetic marker. The maximum-likelihood tree includes reference sequences from the Giant Virus Database (GVDB). Strips showing the site where the MCP was obtained as well as order-level classification of reference sequences are present.

Large phages had similar functional capacities to their smaller counterparts, having AMGs present in a broad range of categories despite there only being 16 identified genomes (Figure S1). The only unique pathway found in these large viruses was thiamine metabolism represented by one AMG, sulfur carrier protein ThiS adenylyltransferase. Besides this pathway, they had a larger proportion of AMGs involved in lysine degradation, peptidoglycan biosynthesis, carbon fixation in photosynthetic organisms, and glutathione metabolism.

When diving into specific AMGs found among the large phages, photosystem II P680 reaction center D1 and D2 proteins were found which protect photosystem II against photoinhibition and UV-B effects (G Schuster, et al, 1988, L sass et al, 2004; C Monod et al. 1994). The CpeT protein, a component of phycoerythrin in cyanobacteria, was also found. Phosphoribulokinase was also found in these large phages, which plays a vital role in photosynthetic enzyme activity modulation (M Rault et al, 1991).

### Phage Host Predictions

The species classification of 329 (16.7%) phages provided some insight into the hosts of the viruses, but most prokaryotic viruses could not be classified. Using a variety of bioinformatic approaches, however, hosts could be predicted for 312 (15.9%) phages. The families of the dominant hosts of phages from both Saint Martin and Cox’s Bazer were Rhodobacteraceae, Flavobacteriaceae, Enterobacteriaceae, and Vibrionaceae (Figure S2). Saint Martin, however, had a much higher proportion of *Vibrio* host predictions than Cox’s Bazar, as well as the less dominant predicted host families of Alteromonadaceae and Chitinophagaceae. Interestingly Cox’s Bazar phages were predicted to infect hosts of the Rickettsiales order, which is a group commonly associated with disease and parasitism in marine organisms. Cox’s Bazar also contains a number of predicted hosts of the Mycobacteriacea family, members of which are known human pathogens involved in diseases like tuberculosis and leprosy, suggestive of wastewater input.

### NCLDV Diversity and Function

In total, 5 NCLDV MAGs (metagenome assembled genomes) were assembled from Cox’s Bazar but none from Saint Martin, although NCLDV contigs were present in Saint Martin as well. Their genome sizes ranged from 83kbp to 876kbp. Four belonged to the order Imitervirales and one belonged to Pandoravirales. GC percentages for these genomes ranged from 25.88-60.9. Maps of the genomes also show a wide number of coding regions with metabolic functions as well as genes involved in DNA processing (Figure 6A). Gene counts from the genomes range from 72-831 and tRNA counts range from 0-10 per genome (Table S1).

**Figure 6.**
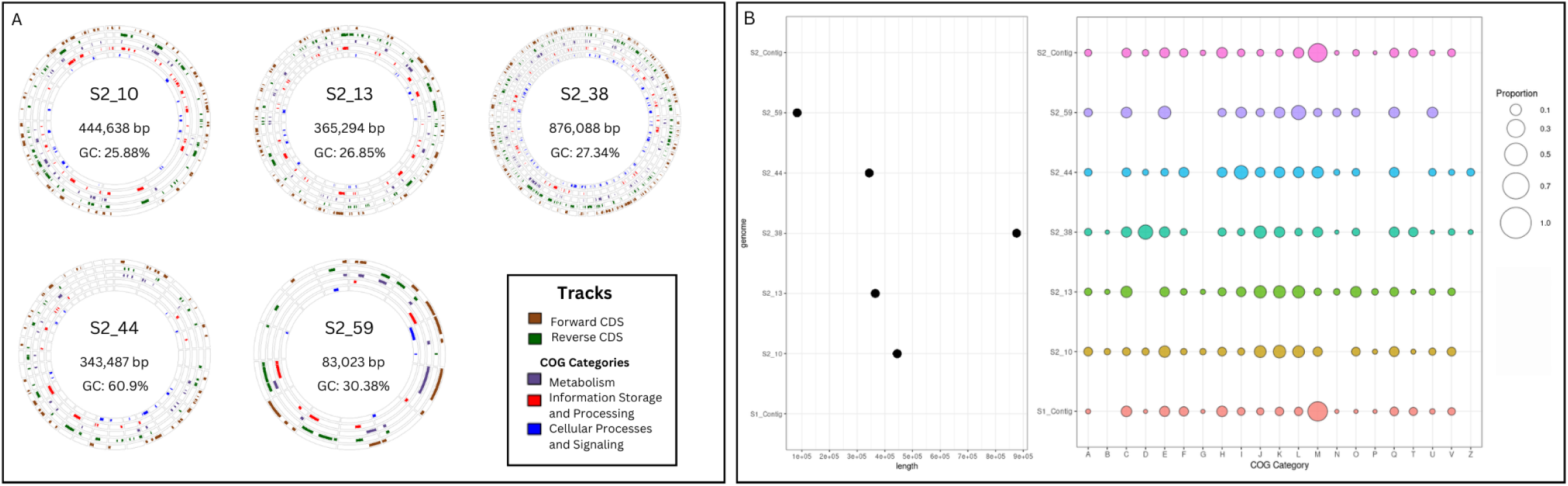
NCLDV Genomes and functional potential. **(A)** Genome plots of the 5 recovered NCLDV genomes, showing GC percentage and total length. Tracks are also shown for forward and reverse coding sequences (CDS) as well as the three major groupings of COG categories (Clusters of orthologous genes): Metabolic, information, and cell processing. **(B)** Functional analysis of the genomes was done by annotating the genomes and unbinned contigs against the GVOG, Pfam, and EggNOG databases using HMMER. What is shown are COG (clusters of orthologous groups) categories and the proportion of genes encoded by each genome or unbinned NCLDV contig in that respective category. COG categories are: (A: RNA processing; B: Chromatin dynamics; C: Energy production; D: Cell cycle control; E: Amino Acid metabolism; F: Nucleotide metabolism; G: Carbohydrate metabolism; H: Coenzyme metabolism; I: Lipid metabolism; J: Translation; K: Transcription; L: Replication and repair; M: Cell membrane biogenesis; N: Cell motility; O: Post-translational modification; P: Inorganic ion transport; Q: Secondary structure; T: Signal transduction; U: Intracellular trafficking; Y: Nuclear structure; Z: Cytoskeleton.

Apart from the 5 NCLDV MAGs we discovered, we also focused on the phylogenetic diversity of the major capsid protein (MCP) that we could detect in NCLDV contigs to get a broader view of NCLDV diversity in this system (Figure 5B). A total of 53 MCPs were recovered from our assembled contigs of which 51 came from Cox’s Bazar and only 2 from Saint Martin, which we compared with reference MCP sequences of cultured NCLDV representatives. The majority of these MCP fell into the Algavirales or Imitervirales order, with one clustering inside the Pandoravirales order. Both MCPs recovered from Saint Martin were found within the Imitervirales order. It is to be noted that multiple copies of MCP are usually present within many NCLDV genomes, and some cases of transfer of this gene across the orders possibly happened - as suggested by the placement of reference genome NCLDVs in our phylogeny. Nevertheless, this gene is frequently used for phylogenetic assessment given its near-universal presence across a broad range of NCLDVs (Moniruzzaman et al., 2017; Moniruzzaman et al., 2016; Larsen et al, 2008). In addition to the 5 MAGs, 39 unbinned NCLDV contigs from Saint Martin and 60 from Cox’s Bazar were recovered and used for functional profiling.

NCLDV genomes from the Bay of Bengal encode a wide range of functional capabilities, with two of the MAGs and 5 unbinned NCLDV contigs from both sites having genes involved in carbohydrate metabolism and transport (Figure 6B). The largest proportion of genes in both sets of unbinned contigs included those involved in cell membrane biogenesis. Two of the giant virus MAGs had proteins for cytoskeleton manipulation which have been described in previous studies (Da Cunha et al., 2022). Three contained proteins for chromatin modification. Although less common, two of the NCLDV MAGs and two unbinned contigs encoded genes for ion transport.

## Discussion

Marine viruses are ubiquitous in the world’s oceans and their diversity and abundance typically reflect that of their hosts whether prokaryotic or eukaryotic (Allen et al., 2017). As viruses can only reproduce by infecting their hosts, they are tightly linked to processes impacting their hosts’ availability. In a sister study to this one, the microbial communities at each of these sites were characterized (Akter et al., 2023), which found that Cox’s Bazar had a higher alpha diversity and species richness in terms of both eukaryotic and prokaryotic communities compared to Saint Martin. Likewise, we found viral diversity was higher at Cox’s Bazar compared to Saint Martin, which suggests that a more diverse host pool could result in a more diverse viral pool. However, studies have shown conflicting results in marine environments and also suggest taxonomic resolution plays a role in this relationship (Gregory et al 2019; Luo et al. 2021). Nevertheless, multiple factors could contribute to an observed higher viral diversity, as well as abundance, at Cox’s Bazar. Cox’s Bazar had on average higher temperatures, lower salinity, and higher total dissolved solids (TDS), which may play a role in viral diversification. Furthermore, Cox’s Bazar’s proximity to a developed and populated coastline could also enhance viral abundance and diversity here – as multiple sources like freshwater inputs from the rivers and wastewater that flow into the bay could introduce viruses in this environment compared to Saint Martin.

Surprisingly, only 61 phage genomes (3.01%) were shared between two sites only ~60 miles away from each other. Despite being planktonic and movement mediated by oceanic currents, it seems that viruses may exhibit high endemicity in this region, a phenomenon that has previously been reported for phages in the ocean’s surface waters (Thurber, 2009; Gregory et al 2019). Because of the fairly similar water temperatures and time of sampling, which are known to impact viral community structure (Aylward et al, 2017; Gregory et al, 2019), other factors could be contributing to the distinctions between the two sites. Furthermore, metagenomes are a snapshot of a community in time, and only two samples were collected at these sites, more would be needed statistically confirm this. Future studies will be needed, that include spatial and temporal sampling schemes, for a comprehensive assessment of the biotic and abiotic drivers of viral community differences in these sites.

One convincing possibility for the striking differences in the viral communities of these sites is that human land usage and water inputs are impacting these sites to different degrees. A study by French et al. (2022) found that in a river environment, urban development and farming had significant impacts on the virus community present in the water. In our study, the influence of land usage and terrestrial water input may explain the detection of phages associated with prokaryotic hosts that cause plant diseases such as the *Ralstonia* phage, which was identified in high abundance at Cox’s Bazar. Furthermore, anthropogenic water inputs to a marine environment typically add nutrients such as phosphorus and nitrogen which are often thought of as limiting nutrients for photosynthetic organisms in the photic zone (Naden et al., 2016). These bloom-forming conditions may have resulted in the observed high abundance of cyanophages and *Celluphaga* phages in Cox’s Bazar site compared to Saint Martin. Cyanophages infect cyanobacteria and the abundance of these phages may reflect an increase in cyanobacterial growth from the nutrient input. Additionally, phages of the *Cellulophaga* species tend to infect members of the *Cellulophaga* bacterial genus. These bacteria mainly consume organic matter from decaying microbial cells (Pati et al., 2011), and they have been detected as enriched at Cox’s Bazar relative to Saint Martin in the sister study by Akter et al. (2023). Although the high abundance of these phages doesn’t always indicate the high abundance of their host, their high abundance of these phages suggests they are active in these environments, considering the often-quick decay rates of phage particles (Noble & Fuhrman, 1997; Suttle & Chen, 1992).

The larger viral abundance and diversity at Cox’s Bazar likely reflects the larger diversity and abundance of AMGs found within the genomes of these viruses. AMGs have been shown to be important factors in modulating host metabolism (Heyerhoff et al., 2022). The presence of a broader range of these AMGs at Cox’s Bazar could imply a broader range of metabolisms used by hosts from which these genes were likely acquired - as AMGs have been shown to vary by host diversity (Luo et al., 2022). Many AMGs are possibly used by viruses to increase virus production by modulating rate-limiting processes in the host (Sullivan, 2006).

The pathway that was the most differentially enriched in Cox’s Bazar compared to Saint Martin was glutathione metabolism, having 5x more genes found in Cox’s Bazar. This pathway has been shown to be implicated in antioxidant defense, nutrient metabolism, and regulation of cellular events (Wu et al., 2004), suggesting phages in Cox’s Bazar may have the ability to manipulate these processes in their hosts. Peptidoglycan biosynthesis is another pathway that is only found in Cox’s Bazar. This pathway has been shown to be utilized by some phages during infection as disrupting peptidoglycan cell walls of bacteria could make viral entry easier (De Smet et al., 2016). The presence of this pathway at only one site could suggest, as it has been hypothesized, that there are phage-specific strategies for infection and metabolic reprogramming of host cells (Howard-Varona et al., 2020).

Jumbo phages have gained recent interest for their large sizes and therapeutic potential, being seen as natural antibacterials against pathogenic coral-infecting bacteria (Jacquemot et al., 2018) as well as their potential to fight human pathogens. These phages have previously been characterized based on their genome size, biogeography, and infection strategies (Weinheimer & Aylward, 2022). Here, we show that there are jumbo phages from multiple distinct phylogenetic groups in the TerL gene phylogeny of phages, with a large clade particularly enriched in these phages. A similar phylogenetic approach can be used to identify and classify novel jumbo phages as well as looking into the evolutionary history of these large phages in other marine environments. This phylogenetic relationship helped to confirm the presence of jumbo phages at Cox’s Bazar through phylogenetic similarity to known jumbo phages. Jumbo phages are a part of the larger and more diverse viral community at Cox’s Bazar, potentially due to an increased amount of nutrients to support larger host populations. It is important to iterate that recovery of a large number of jumbo phage-specific phylogenetic markers in our study is consistent with the observations that large viruses are typically enriched within the cellular size fraction. Thus, our study re-affirms the importance of investigating the cellular size fraction for the assessment of viral diversity along with the < 0.22-micrometer fraction that is routinely used (Palermo et al., 2021).

Large phages in our study contained a large number of AMGs relative to the small number of identified genomes of 16. Due to their large genome sizes, these phages have a larger capacity to encode AMGs and confer metabolic changes to their host. Among large phage-encoded metabolic potentials from the genomes in this study, we found a high number of AMGs involved in the control of photosynthesis such as photosystem II P680 reaction center D1 and D2 proteins. These genes have been shown to be functional in the host and have been hypothesized to increase phage fitness through supplementing and protecting the host’s photosynthetic machinery (Lindell, 2005; Ban 2019). Jumbo phages remain a poorly studied group of phages and further investigations into their full ecological potential are required.

Viral host prediction can be a challenging task when culture-confirmed interactions are not available from the study environment. Our host prediction approach leveraging multiple bioinformatic evidence revealed a large proportion of host-virus interactions within the Rhodobacteraceae. This result is consistent with the fact that Rhodobacteraceae is highly abundant in coastal waters across the world, playing a role in organic matter degradation (He et al., 2017). Interestingly, Cox’s Bazar had host predictions to members of the Rickettsiales order which is commonly associated with disease and parasitism in marine organisms. Cox’s Bazar also contains a number of hosts predicted from the Mycobacteriaceae family which is known for causing diseases such as tuberculosis and leprosy in humans (Böttger 1991). These pathogenic hosts are likely the result of the many anthropogenic influences in the Cox’s Bazar waterways. Microbial analysis of Cox’s Bazar also suggested that these waters could be harboring pathogenic organisms due to the significant number of identified virulence genes (Akter et al, 2023).

While only Cox’s Bazar samples had a high enough abundance of NCLDVs to recover MAGs, NCLDV contigs were still found in Saint Martin. The majority of MCPs found in the data are from the orders Algavirales and Imitervirales, which are the two largest orders of NCLDVs and are generally the most abundant in coastal waters (Ha et al., 2023). The large genomes of NCLDVs have been shown to greatly alter host metabolism as they contain many genes that mimic host homologs and can hijack host machinery to turn the host’s metabolism towards making viruses (Moniruzzaman et al., 2020). Many of these genes are unique to NCLDVs in the virus community and are likely acquired from cellular hosts (Fixen et al., 2022). Functional annotation of our NCLDV genomes revealed genes such as those involved in DNA packaging (histones), cytoskeletal manipulation, and ion transport. It is hypothesized that NCLDV histones mimic host histones to protect viral DNA from degradation, forming similar structures to nucleosomes (Liu et al, 2021). Ion transporters have long been found in viruses and likely play a role in viral entry through depolarization of the membrane, and nutrient acquisition to provide the host with additional nutrients to support viral replication (Monier et al., 2017). Cytoskeletal manipulation proteins have been suggested to be used by NCLDVs to hijack the host’s vesicular trafficking, bringing cargo to the viral factory for viral assembly and production (Arantes et al., 2016; Wilson et al., 2005). Our data on the NCLDVs represents the first report of NCLDVs in the Bay of Bengal - highlighting their potential role in the microbial food web in this region and adds to the growing body of knowledge on the diversity and genomic potential of NCLDVs in the world’s ocean.

Despite the valuable insights gained from this study, it is important to acknowledge and address several limitations that may have influenced the findings. The limited number of sites by which to compare viral populations limits the scope of this study, as only two sites were selected in this large bay. The pooling of the two samples from these two sites also limits our ability to evaluate drivers of the viral community at a finer spatial scale. Additionally, more physicochemical parameters would have been helpful in order to provide further evidence for anthropogenic influences and support hypotheses regarding the differences in the viral communities. However, the primary motivation for this study was to evaluate the diversity of viruses and their hosts in this understudied, yet highly important ecosystem. Future analyses with more samples collected at a higher spatial resolution will be necessary to discover the drivers of virus community structure and dynamics. The current repertoire to classify and identify viral hosts is also limited, and we could classify and identify host affiliation of only a small proportion of the viruses. In spite of these limitations, this study provides the necessary foundation for future research on the microbial and viral communities in the BoB.

## Conclusion

Our study aimed to address the knowledge gap regarding the viral community diversity and structure within an understudied ecosystem, the Bay of Bengal (BoB). Given the region’s susceptibility to frequent monsoons, flooding, and vulnerability to sea-level rise, the BoB holds research significance as its neighboring nations grapple with these challenges. Considering the crucial role of microbial populations in supporting fisheries that sustain numerous livelihoods, our research on a portion of the BoB’s virus population contributes to the broader understanding of the factors that constrain microbial population dynamics and their contribution to the food web in this ecosystem. Our study underscores the importance of continued research in comprehending the intricate dynamics and ecological significance of microbial populations in sustaining the blue economy, as well as the anthropogenic influences affecting viral and microbial dynamics and abundance in this region. In addition, the detection of giant viruses and large phages in the Bay of Bengal underscores the need to extend marine virological research into less-studied ecosystems, which will enhance our comprehension of the rich diversity of marine viruses and better elucidate their evolutionary histories. Finally, this work will be a crucial reference for future research on deciphering the viral impact on microbial food web dynamics in the Bay of Bengal.

## Data Availability

Raw sequence reads are available at the NCBI database under the bioproject number PRJNA936489. Genomes of NCLDVs and large phages, as well as the information used to make phylogenetic trees, functional annotations, and code used to make the plots can be obtained from (https://figshare.com/projects/Phylogenetic_diversity_and_functional_potential_of_large_and_cell-associated_viruses_in_the_Bay_of_Bengal/171777). Bootstrapped versions of maximum likelihood TerL and MCP trees in newick format are also provided in the same repository. The pipeline for identification of Giant virus Genomes (PIGv) can be found at https://github.com/BenMinch/PIGv.

## Supporting information

Supplementary_info_figures_and_tables_Minch_et_al_2023

## Acknowledgement

The work was supported by funding from Rosenstiel School of Marine, Atmospheric and Earth Sciences, University of Miami to MM.

## Author Contributions

BM performed the majority of the data analysis, interpreted results, visualized figures and drafted the original manuscript. AW, SA and MSR performed data analysis and critically reviewed the manuscript. MFA, SRRA and MAKP aided in sample collection, provided laboratory and reagent support, and critically reviewed the manuscript. MM conceived the study, provided funding and supervised the research.

## Figure Descriptions

**Figure S1. Large phage metabolic potential.** A barplot showing auxiliary metabolic genes found in the binned and unbinned large phage genomes using VIBRANT. Colors represent broader metabolic pathways.

**Figure S2. Phage Virus-Host Prediction.** Host prediction was performed using iPHOP and represented here is the predicted number of virus-host pairs to the classification level of host family. In total 312 Host-Virus pairs were predicted using this method which represents about 16% of our total viral populations identified.

**Table S1. NCLDV Genome Overview.** Information about recovered genomes including size, classification, GC percentage, gene count, coding percentage, NCLDV marker genes, and tRNAs.

