## Supplementary_info_figures_and_tables_Minch_et_al_2023 for "Phylogenetic diversity and functional potential of large and cell-associated viruses in the Bay of Bengal"

### This file includes:

Supplementary figures S1, S2 and supplementary table S1

**Figure S1. Jumbo phage metabolic potential.** A barplot showing auxiliary metabolic genes found in the binned and unbinned jumbo phage genomes using VIBRANT. Colors represent broader metabolic pathways.

**Figure S2. Prokaryotic Virus-Host Prediction.** Host prediction was performed using iPHOP and represented here is the predicted number of virus-host pairs to the classification level of host family. In total 312 Host-Virus pairs were predicted using this method which represents about 16% of our total viral populations identified.

**Table S1. NCLDV Genome Overview.** Information about recovered genomes including size, classification, GC percentage, gene count, coding percentage, NCLDV marker genes, and tRNAs.

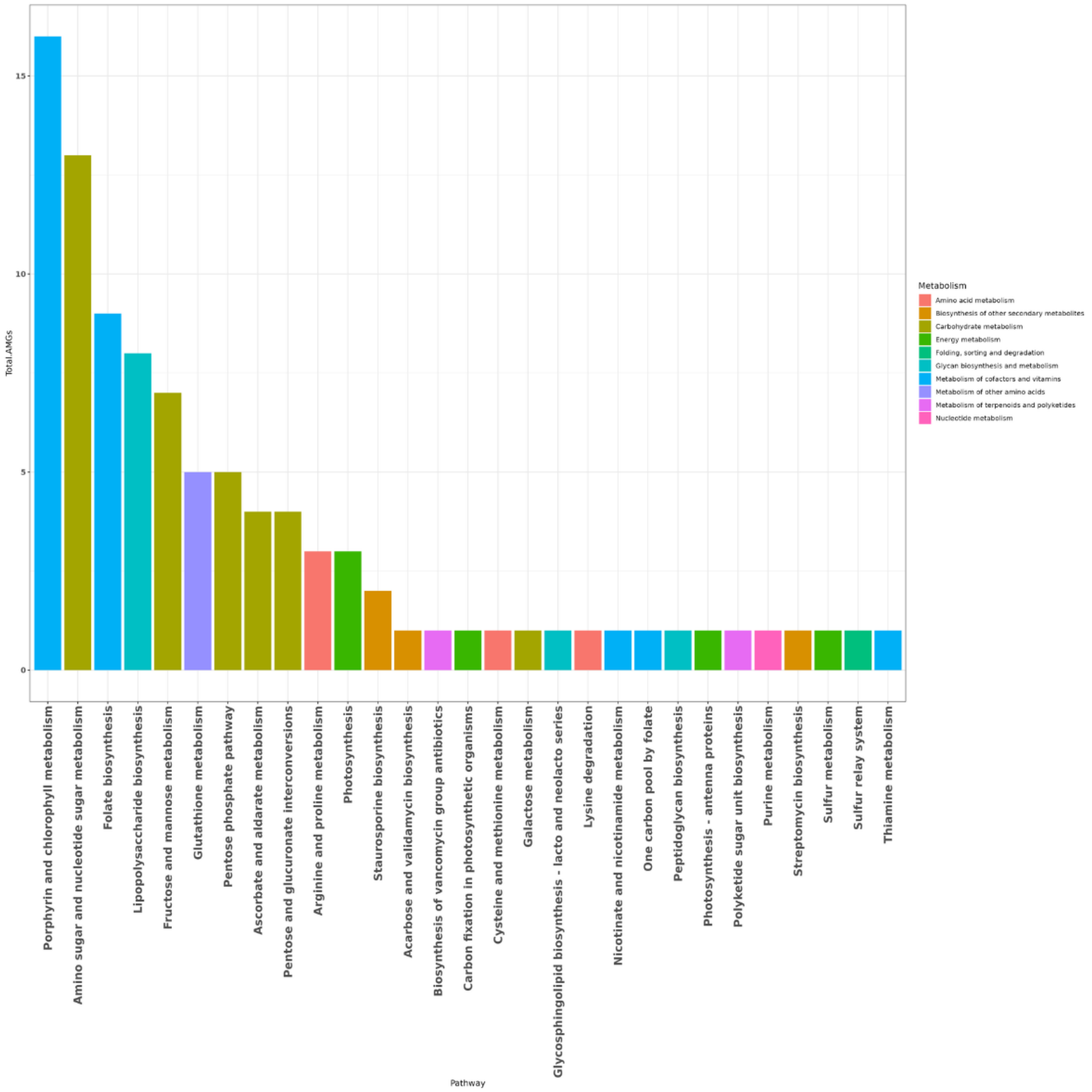


**Figure S1. Jumbo phage metabolic potential.** A barplot showing auxiliary metabolic genes found in the binned and unbinned jumbo phage genomes using VIBRANT. Colors represent broader metabolic pathways.

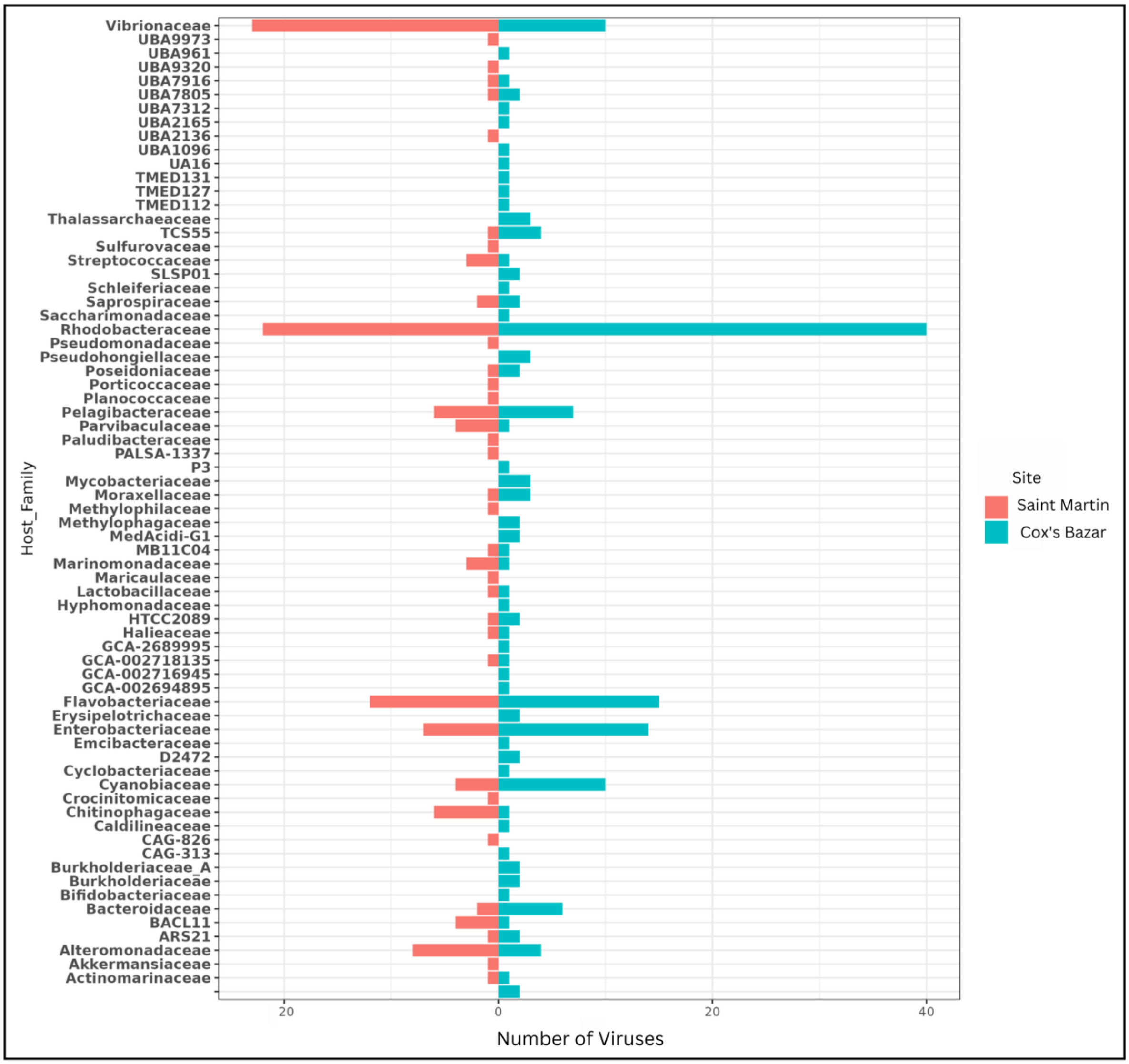


**Figure S2. Prokaryotic Virus-Host Prediction.** Host prediction was performed using iPHOP and represented here is the predicted number of virus-host pairs to the classification level of host family. In total 312 Host-Virus pairs were predicted using this method which represents about 16% of our total viral populations identified.

| Genome | Size (bp) | GC % | Gene Count | Coding % | tRNAs | NCLDV Marker Genes | Family | Order |
|---|---|---|---|---|---|---|---|---|
| S2_10 | 444638 | 25.88 | 290 | 93.53 | 10 | 9 | Mesomimiviridae | Imitervirales |
| S2_59 | 83023 | 30.38 | 72 | 90.83 | 1 | 4 | Mesomimiviridae | Imitervirales |
| S2_13 | 365294 | 26.85 | 313 | 91.52 | 9 | 8 | Mesomimiviridae | Imitervirales |
| S2_38 | 876088 | 27.34 | 831 | 92.85 | 0 | 8 | Mimiviridae | Imitervirales |
| S2_44 | 343487 | 60.9 | 294 | 87.94 | 0 | 4 | incertae_sedis | Pandoravirales |

**Table S1. NCLDV Genome Overview.** Information about recovered genomes including size, classification, GC percentage, gene count, coding percentage, NCLDV marker genes, and tRNAs.
